# Establishing an *ex vivo* porcine skin model to investigate the effects of broad spectrum antiseptic on viable skin microbial communities

**DOI:** 10.1101/2025.04.03.647100

**Authors:** E.C. Townsend, K. Xu, K. De La Cruz, L. Huang, S. Sandstrom, J.J. Meudt, D. Shanmuganayagam, A. Huttenlocher, A.L.F. Gibson, L.R. Kalan

## Abstract

Incomplete antiseptic efficacy against potentially pathogenic microbial taxa places some patients at disproportionate risk for developing a surgical site infection. Laboratory models capable of interrogating the effects of antiseptics on the skin and its complex microbial communities are desperately needed to improve and better tailor antiseptic formulations. This work aims to establish an *ex vivo* porcine skin model to explore the impact of topical antiseptics on complex skin microbial communities and superficial skin lipids. Microbiome samples were treated with propidium-monoazide to selectively evaluate DNA from viable microorganisms. Bacterial abundances were assessed via viability-qPCR and quantitative culture. Viable community populations were evaluated with 16S rRNA gene sequencing. Epidermal biopsies were collected at multiple timepoints for lipidomic assessment via LC/MS. The *ex vivo* environment promoted shifts in porcine skin lipid composition and microbial communities over the experiment’s duration. Compared to water treated control skin, skin treated with the antiseptic chlorhexidine gluconate had significantly lower culturable and bioburden determined by viability-qPCR. Compared to water treated skin, viable microbial communities on CHG treated skin displayed greater relative abundance of several gut associated and Gram-negative bacterial taxa, including *SMB53, Turicibacter, Pseudomonas* and *Proteus*. Collectively these findings highlight the utility of an ex vivo porcine skin system for interrogating the impacts of antimicrobial disruption to complex microbial ecosystems, and ultimately for the future testing and development of improved antiseptic formulations.

## Introduction

Surgical site infections (SSI) pose a substantial burden to affected patients and the healthcare system.^1–4^ Pre-surgical antiseptics intentionally and effectively reduce microbial bioburden at the time of surgery to help prevent SSI from developing. However, contrary to popular belief, complete sterility is not achieved.^5^ Rather, antiseptics temporarily disrupt skin microbial communities and can promote enrichment of several potentially pathogenic taxa such as *Pseudomonas, Escherichia, Acinetobacter*, and *Bacillus*.^5^ Incomplete antiseptic efficacy against these pathogenic Gram-negative and biofilm forming species places some patients at disproportionate risk for developing an SSI.^1,6,7^ To better characterize the current limitations of antiseptics and ultimately improve antiseptic formulations, comprehensive laboratory models to evaluate the impacts of antiseptics on the skin and its complex microbial communities are needed.

Model systems to study skin physiology, immunology, and wound healing include animal models (e.g. mouse, porcine), Reconstructed Human Epidermis, and *ex vivo* human or animal skin tissue.^8–10^ Of these, *ex vivo* porcine skin tissue may offer the greatest opportunity for interrogating the complex ecosystems of bacteria, fungi, viruses, and archaea that comprise the skin microbiome and how these communities respond to acute disruptions like antiseptic exposure. E*x vivo* skin contains all the microstructures (e.g. hair follicles, sweat glands, sebaceous glands) that support skin microbial communities.^8^ Porcine and human skin are highly similar in structure, biochemistry, as well as keratinocyte and resident immune cell functions,^11–14^ making porcine skin a valuable model for studying human skin function and wound healing.^11,14–16^ Microbial communities that reside on porcine and human skin also overlap, containing 97% of the same bacterial genera.^17,18^ Unlike *ex vivo* human skin tissue, which is acquired following surgical removal, porcine skin is not always treated with antiseptics prior to removal from a euthanized animal and can be acquired with its skin microbial communities intact.^18^

Chlorhexidine gluconate (CHG) is one of the preferred surgical antisepsis due to its ability to reduce culturable microbial bioburden for up to 48 hours post-application.^19–22^ Prior to surgery, many surgical centers have patients shower with 4% CHG soap the night before and morning of their surgery. Immediately prior to incision CHG is then applied to the surgical site.^22–25^ However, due to CHG’s ability to bind persistent bacterial DNA to the surface of the skin,^26,27^ sequencing-based efforts to quantify and characterize the impact of CHG on the skin microbiome have yielded inconsistent results.^26,28–31^ To circumvent this, we optimized a propidium monoazide (PMAxx) based viability assay for selective evaluation of live microorganisms in the skin microbiome pre-and post-antiseptic exposure.^5^ With this method we show that pre-operative CHG effectively reduces viable microbial bioburden but can select for potentially pathogenic taxa, particularly Gram-negative and biofilm forming bacteria. This underscores the need to develop laboratory models for the study and improvement of antiseptic formulations. To address this, we utilized this viability assay to characterize the effects of CHG on skin microbial communities in this *ex vivo* porcine skin system.

In this work we establish an *ex vivo* porcine skin model to explore the impact of CHG antiseptic on the skin microbiome. Consistent with our studies in human surgical populations, this work confirms that application of CHG effectively reduces viable microbial bioburden. However, this effect is temporary and accompanied by the enrichment of several potentially pathogenic taxa. We show that the laboratory model resembles a moist skin environment, and the skin undergoes lipid remodeling. Collectively, these findings highlight the advantages of an *ex vivo* porcine skin system for interrogating the impacts of topically applied antimicrobials and other chemical or mechanical disruptions to the skin microbiome.

## Methods

### Ex vivo porcine skin tissue handling

*Ex vivo* porcine skin tissue was obtained from 6 month to 2 year old Wisconsin Miniature Swine (WMS)™ that were bred and maintained at the University of Wisconsin - Madison. All tissue came from WMS that had been utilized and euthanized as part of other research studies being conducted that did not involve antiseptic exposure to the dorsal skin, antibiotic treatment, or induced immunocompromise. *Ex vivo* tissue was washed with sterile water and sterile gauze to remove superficial dirt and blood. Hair was removed by trimming the hair with scissors followed by shaving with a 2 or 3 blade razor. Tissue was again rinsed with sterile water and gauze until clean. Subcutaneous muscle and fat were removed and subcutis tissue was trimmed with a scalpel to as even as a depth as possible (approximately 1 cm thick). Outer edges of the tissue, roughly 1 cm, were removed. Tissue was then divided into roughly equal, 4×7 inch sections, one for each experimental group. For longer experiments extending over several days, tissue sections were placed on 9×9 inch Dulbecco’s Modified Eagle Medium (DMEM) gel plates. These plates were made by adding 76.5% high glucose DMEM (Cytivia, Marlborough, MA) + 8.5% Fetal Bovine Serum (FBS; Thermo Fisher Scientific, Waltham, MA) + 15% of 2-percent agarose in 1X phosphate buffered saline (PBS) by volume. Tissue was stored on DMEM gel plates at 37°C in a 5% CO_2_ incubator with daily plate changes for the duration of the experiment. Images displaying the *ex vivo* model are in **Supplemental Figure 1**.

### Experimental Design

To explore the effects of CHG antiseptic on the *ex vivo* porcine skin epidermal lipid composition and microbiome, sections of *ex vivo porcine* skin from 3 porcine donors were either i) bathed twice with 4% CHG soap twelve hours apart followed by application of 2% CHG in 70% Isopropanol (IPA) to mimic standard pre-surgical preparations (in orange, **Fig 1A**); or ii) bathed twice with sterile water twelve hours apart followed by a single application of 2% CHG in 70% IPA to mimic local antiseptic application (e.g. to mimic skin preparation before a blood draw; yellow); or iii) bathed three times with sterile water as a control (teal-green). Swabs of the skin microbiome were taken at baseline, immediately after (0 hours) as well as 6, 12, 24 and 48 hours after the final treatment intervention. Microbial burden was then measured with quantitative microbial culture and viability-qPCR of the bacterial 16S rRNA gene (detailed below). To investigate CHG’s potential effects on epidermal lipid and short chain fatty acid profiles, punch biopsies of the epidermis were taken from two of the porcine donors in this experiment (detailed below).

**Figure 1:**
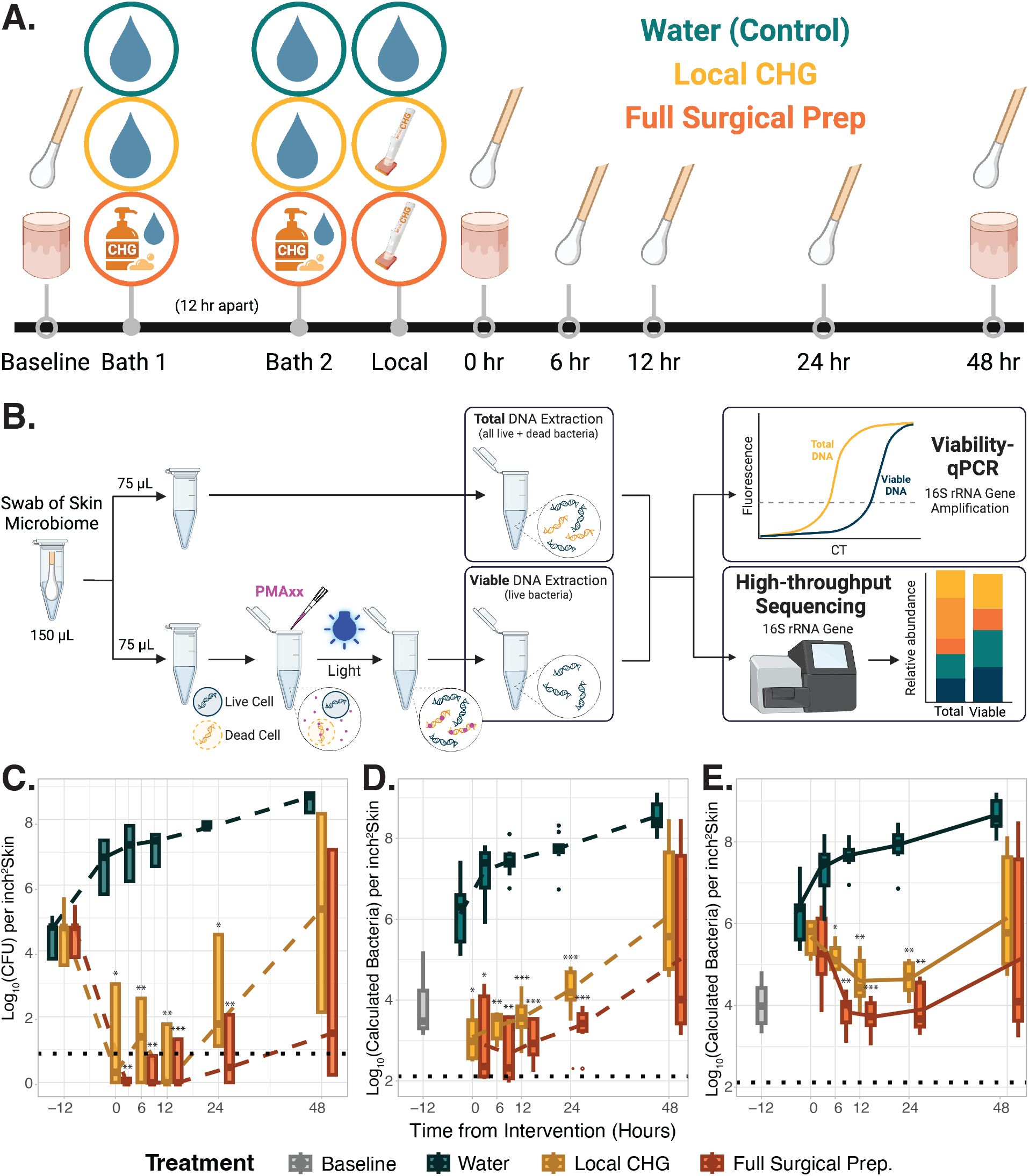
Chlorhexidine gluconate antiseptic reduces viable microbial bioburden. **A**.Experimental design. Sections of *ex vivo* porcine skin from 3 porcine donors were treated i) bathed twice with 4% CHG soap twelve hours apart followed by application of 2% CHG in 70% Isopropanol (IPA) to mimic standard pre-surgical preparations (bottom, orange); ii) bathed twice with sterile water twelve hours apart followed by a single application of 2% CHG in 70% IPA to mimic local antiseptic application (middle; yellow); or iii) bathed three times with sterile water as a control (top; teal-green). Swabs of the skin microbiome were taken at baseline, immediately after (0 hours) as well as 6, 12, 24 and 48 hours after the final treatment intervention. Punch biopsies of the epidermis from 2 of the porcine donors were taken at baseline, immediately after treatment intervention as well as the endpoint for lipidomic assessment. **B**. Diagram of sample processing for viability q-PCR and sequencing. Skin microbiome samples were split. Half of each sample was treated with PMAxx to selectively amplify DNA from viable bacteria. The remaining half of the sample remained untreated to evaluate the DNA from both live and dead bacteria (total DNA). Total and viable bioburden was evaluated via quantitative PCR of the bacterial 16S Ribosomal RNA gene (viability-qPCR). Viable and total microbial community composition were then evaluated via high-throughput 16S rRNA gene sequencing. **C**. Viable microbial bioburden per inch^2^ skin as determined by quantitative bacterial culture for each of the experimental groups in panel A over time. **D**. Viable microbial bioburden per inch^2^ skin as determined by viability-qPCR for each of the experimental groups over time **E**. Total microbial bioburden per inch^2^ skin measured by viability-qPCR for each of the experimental groups over time. For panels **C-E**, bioburden in the local CHG and full surgical CHG preparation groups were compared to the water treated control group at each respective time point via t-tests with welches correction. For each panel black dashed lines indicate the lower limit of each assay’s detection. * indicates p-value < 0.05. ** indicates p-value < 0.01. *** indicates p-value < 0.001.

### Sample Collection

Swabs of the skin microbiome were collected from a 1 inch^2^ area of skin via the Levine technique.^32^ Swabs designated for DNA extraction were placed into 155 μl of 1% Bovine Serum Albumin (BSA) in 1x PBS and were stored at 4°C for less than 30 minutes before processing selective detection of DNA from viable organisms (per below) and eventual DNA extraction. Swabs designated for microbial culture were also taken using Levine’s technique from a 1 inch^2^ area of skin into 100 µl of 1% BSA in 1x PBS and were stored at 4°C for less than 2 hours before being processed for microbial culture.

### Quantitative Microbial Culture

Swabs designated for microbial culture were spun down using DNA IQ Spin Baskets (Promega, Madison, WI). Samples were serially diluted with 1X PBS and plated onto Tryptic Soy Agar (TSA) then incubated at 35°C overnight for quantitative bacterial culture.

### Selective Detection of DNA from Viable Microorganisms

Swabs collected into PBS + 1% BSA were spun down using DNA IQ Spin Baskets (Promega, Madison, WI) and each sample was split into two equal 75 µl portions. To quantify and sequence the DNA from all (live and dead) members of the *ex vivo* skin microbial communities, one portion of each sample was placed directly into −20°C storage. To selectively quantify and sequence DNA from only live microbes within skin microbial communities, the other portion of each sample was processed with a modified propidium monoazide (PMAxx, Biotium, Fremont, CA) based viability assay (**Fig. 1B**). In short, PMAxx irreversibly binds to free DNA and DNA within permeable (dead) cells, allowing for selective amplification and sequencing of only the non-PMAxx-bound DNA within intact (viable) cells. All steps involving PMAxx were done in a dark room. PMAxx was added to achieve a final concentration of 10 µM in each sample. Samples were rocked at room temp for 10 minutes, exposed to blue light for 15 minutes via the PMA-Lite LED Photolysis Device (Biotium, Fremont, CA), then spun at 5000 g for 10 minutes. Both the PMAxx treated portion and untreated portion of each sample were stored at −20°C before DNA extraction.

Initial experiments aimed to optimize the viability assay parameters for selective evaluation of viable microbes within the complex microbial communities residing on *ex vivo* porcine skin both in control skin conditions and following application of CHG antiseptic. Swabs of the skin microbiome were collected from sections of *ex vivo* porcine skin. One sample served as a control to determine the anticipated amount of viable bacteria on the skin in normal circumstances, another microbiome sample was boiled at 95°C for 10 minutes to heat-kill the majority of the bacteria present, as a negative control. A third sample was taken from a section of skin after application of 2% CHG in 70% isopropanol antiseptic. Samples were split, with half treated with PMAxx and half remaining untreated and bacterial bioburden determined by viability-qPCR of the 16S rRNA gene (**Fig. 1B**). The number of bacteria in the viable sample portion was divided by the number of bacteria in the total (live + dead) sample portion to determine the percentage of live bacteria within the sample. Each experimental condition was done in triplicate biologic, *ex vivo*, replicates where three technical replicates that were averaged.

### DNA/RNA Extraction, DNA quantification, Library Construction, Sequencing

DNA extraction was performed as previously described with minor modifications.^5^ Briefly, 300 μl of yeast cell lysis solution (from Epicentre MasterPure Yeast DNA Purification kit), 0.3 μl of 31,500 U/μl ReadyLyse Lysozyme solution (Epicentre, Lucigen, Middleton, WI), 5 μl of 1 mg/ml mutanolysin (M9901, Sigma-Aldrich, St. Louis, MO), and 1.5 μl of 5 mg/ml lysostaphin (L7386, Sigma-Aldrich, St. Louis, MO) was added to 150 μl of swab liquid before incubation for one hour at 37°C with shaking. Samples were transferred to a 2 ml tube with 0.5 mm glass beads (Qiagen, Germantown, Maryland) and bead beat for 10 min at maximum speed on a Vortex-Genie 2 (Scientific Industries, Bohemia, NY), followed by a 30 min incubation at 65°C with shaking, 5 min incubation on ice. The sample was spun down at 10,000 rcf for 1 min and the supernatant was added to 150 μl of protein precipitation reagent (Epicentre, Lucigen, Middleton, WI) and vortexed for 10s. Samples were spun down at maximum speed (∼21,000 rcf) and allowed to incubate at RT for 5 min. The resulting supernatant was mixed with 500 μl isopropanol and applied to a column from the PureLink Genomic DNA Mini Kit (Invitrogen, Waltham, MA) for DNA purification using the recommended protocol.

Viability quantitative polymerase chain reaction (viability-qPCR) was performed, to determine the amount of DNA from viable bacteria (treated with PMAxx) and total DNA from both live and dead bacteria (non-PMAxx treated) portion of each sample. In short, 1 µl of extracted DNA was added to a reaction mix containing 5 µl TaqMan Fast Advanced 2X Master Mix (Applied Biosystems, Waltham, MA), 0.5 µl TaqPman 16S 20X Gene Expression Assay with FAM (Applied Biosystems), and 3.5 µl PCR pure water. Samples were run for 40 thermos-cycles on the QuantStudio 7 Flex Real-Time PCR System (Applied Biosystems). Sample DNA concentrations were determined based on a standard curve of 0.015 to 15000 pg/µl DNA extracted from Escherichia Coli (ATCC 1496).

16S rRNA gene V3-V4 region amplicon libraries were constructed using a dual-indexing method at the University of Wisconsin Biotechnology Center and sequenced on a MiSeq with a 2×300 bp run format (Illumina, San Diego, CA). Reagent-only negative controls were carried through the DNA extraction and sequencing process. A 20-Strain Staggered Mix Genomic Material (ATCC, Manassas, VA) served as a positive sequencing control.

### Sequence Analysis

The QIIME2^33^ environment was used to process DNA-based 16S rRNA gene amplicon data. Paired end reads were trimmed, quality filtered, and merged into amplicon sequence variants (ASVs) using DADA2. Taxonomy was assigned to ASVs using a naive Bayes classifier pre-trained on full length 16S rRNA gene 99% OTU reference sequences from the GreenGenes database (version 13_8). Using the qiime2R package, data was imported into RStudio (version 1.4.1106) running R (version 4.1.0) for further analysis using the phyloseq package.^34^ Negative DNA extraction and sequencing controls were evaluated based on absolute read count and ASV distribution in true samples. Abundances were normalized proportionally to total reads per sample. Data was imported into RStudio running R (version 4.2.1) for analysis. Plots were produced using the ggplot2 package. Taxa below 2% relative abundance were pooled into an “Other” category for the relative abundance plots. Bray-Curtis beta diversity metric was utilized to compare sample microbial community structures and all associated plots were ordinated via Non-metric Multidimensional Scaling (NMDS). Univariate and/or multivariate permutational multivariate analysis of variance (PERMANOVA) were used to evaluate associations between microbial community compositions and various experimental groups. Each PERMANOVA was run considering the marginal effects of terms with 9999 permutations using Adonis2 in the vegan r package.^35^ Distances from a groups centroid were calculated using the vegan^35^ betadisper function to evaluate the variability of microbial community compositions of samples from water treated or CHG treated skin. Tukey multiple comparisons of means was then used to determine if the degree of variability within groups were significantly different. MAASLIN2^36^ was utilized to identify significant differences in taxa abundance between various groups and significant correlations of taxa relative abundance over the time. All MAASLIN2 assessments incorporated porcine donor as a random effect.

### Lipidomics

To test for potential effects of the *ex vivo* environment and CHG exposure on the epidermal and sebaceous lipids of *ex vivo* skin, 12mm punch biopsies were collected in triplicate from two of the three experimental replicates (pigs 2 and 3) at multiple timepoints (**Fig. 1A**). Epidermis was excised from each biopsy (**Fig. S1C**), placed directly into a Eppendorf tube and flash frozen in liquid nitrogen for 10-30 seconds, then placed immediately into a −80C freezer for storage prior to lipidomic assessment. Aliquots of the CHG solutions were also provided as controls. Lipid extraction with methyl-tert-butyl ether^37^ and lipidomics were performed at the University of Wisconsin Biotechnology Center.^38^Lipids were identified and quantified via LC/MS/MS and LC/MS respectively using both positive and negative ion modes on the Agilent 1290 Infinity II ultra-high performance liquid chromatography (UHPLC) and Aglilent 6546 QTOF Mass spectrometer (Agilent, Santa Clara, CA). Initial data processing and lipid assignment was done primarily with the Agile acquisition software (Agilent). Due to a bug in Agile software, data from the positive ion mode for pig 3 samples was processed with MS-DIAL.^39^ Lipid quantities in each sample were normalized by tissue mass. Results were imported into R Studio running R (version 4.2.1) for analysis. Plots were produced using the ggplot2 package. To reduce the dimensionality of the lipidomic data and explore the variability in sample lipid composition, Principle Coordinate Analysis (PCA) was conducted via the MixOmics R package.^40^ Univariate and/or multivariate PERMANOVAs were used to evaluate associations between lipid compositions and various experimental groups. PERMANOVAs were all run considering the marginal effects of terms with 9999 permutations using Adonis2 in the vegan r package.^35^ Differential lipid abundance between groups was assessed via DEqMS.^41^ Lipids were considered to be differentially abundant in a given group if they displayed log_2_(fold change) > 2 and adjusted Limma p-value < 0.01.

### Statistical Analyses

Most statistical analyses were conducted in R studio running R (version 4.2.1). Differences between culturable as well as viability-qPCR viable and total microbial bioburden in different experimental groups were analyzed via Prism (version 9.2.0).

### Data Availability

Sequence reads for this project can be found under NCBI BioProject PRJNA1093136. Code for analysis and generation of figures can be found on GitHub at https://github.com/Kalan-Lab/Townsend_etal_ExVivoPorcineSkinCHG.

## Results

### Selective evaluation of DNA from live skin microorganisms

To optimize viability-qPCR parameters for selective quantification of viable microbes, we determined that PMAxx concentrations of 10 µM and 25 µM yielded the most accurate quantification of live bacteria in the control ex vivo porcine skin sample (28±7% and 18±2%, respectively) compared to a heat killed control (2±2% and 3±3%) or from skin treated with CHG antiseptic (10±1% and 9±4%; all p-values < 0.05 heat killed or CHG vs. the respective control, t-tests with Welch’s correction **Fig S2, Table S1**). To maintain consistency with the parameters optimized for human skin microbiome samples,^5^ we proceeded with 10 µM PMAxx for the remaining experiments. Consistent with our observations from human skin microbiome samples,^5^ the comparatively low percentage of live bacteria in the control sample treated with 50 µM PMAxx (7±1%) suggests that excessive PMAxx is cytotoxic.

### Chlorhexidine gluconate antiseptic reduces viable skin microbial bioburden

To explore the effects of CHG antiseptic on the *ex vivo* porcine skin microbiome, sections of *ex vivo porcine* skin from three animal donors received either i) three CHG treatments to mimic standard pre-surgical preparations (orange, **Fig. 1A**); ii) a single application of CHG in IPA to mimic local antiseptic application (yellow); or iii) bathed three times with sterile water (teal-green). At baseline, *ex vivo* porcine skin contained roughly 10^4^ viable bacteria/ inch^2^ determined by quantitative microbial culture and viability PCR (**Fig. 1B-D, Table S2**). Over the course of the experiment, viable microbial bioburden increased to just under 10^9^ bacteria/ inch^2^ *ex vivo* skin in the water treated control group.

Application of CHG induced an immediate reduction in culturable bioburden (**Fig. 1C**). Viability q-PCR revealed that despite negative cultures, approximately 10^3^ viable bacteria/ inch^2^ remained *ex vivo* skin immediately following both the single CHG application and the full surgical prep (**Fig. 1D**). Microbial burden remained lower in the CHG treatment groups over the time course (all p-values < 0.05, t-tests with welches correction, **Fig. 1C-D, Table S2**). Immediately after CHG application, we see no difference in total microbial bioburden (live + dead bacterial DNA) between CHG treated groups and the water control (**Fig 1E, Table S2**). This corroborates previous reports that CHG application leads to persistence of free DNA from lysed cells on the skin. Then a slow decline in total DNA in the antiseptic treated groups occurs over the first 12 hours, suggesting that this free DNA is likely slowly degraded for re-use by the remaining, propagating, members within the community.

### Total and viable microbial community compositions

16S rRNA gene sequencing was used to characterize the impact of CHG antiseptic on viable skin microbial community structure (**Fig. 1A**). At baseline, *Macrococcus, SMB53, Staphylococcus, Streptococcus*, and *Turicibacter* were the dominant taxa within viable and total (both live + dead bacterial DNA) communities (**Fig. 2**). Of the genera identified, 85% have also been found on human skin (**Table S3**). When evaluating all samples together, porcine donor was the largest driver of microbial community composition (univariate PERMANOVA p-value < 0.0001, **Table S4**).

**Figure 2:**
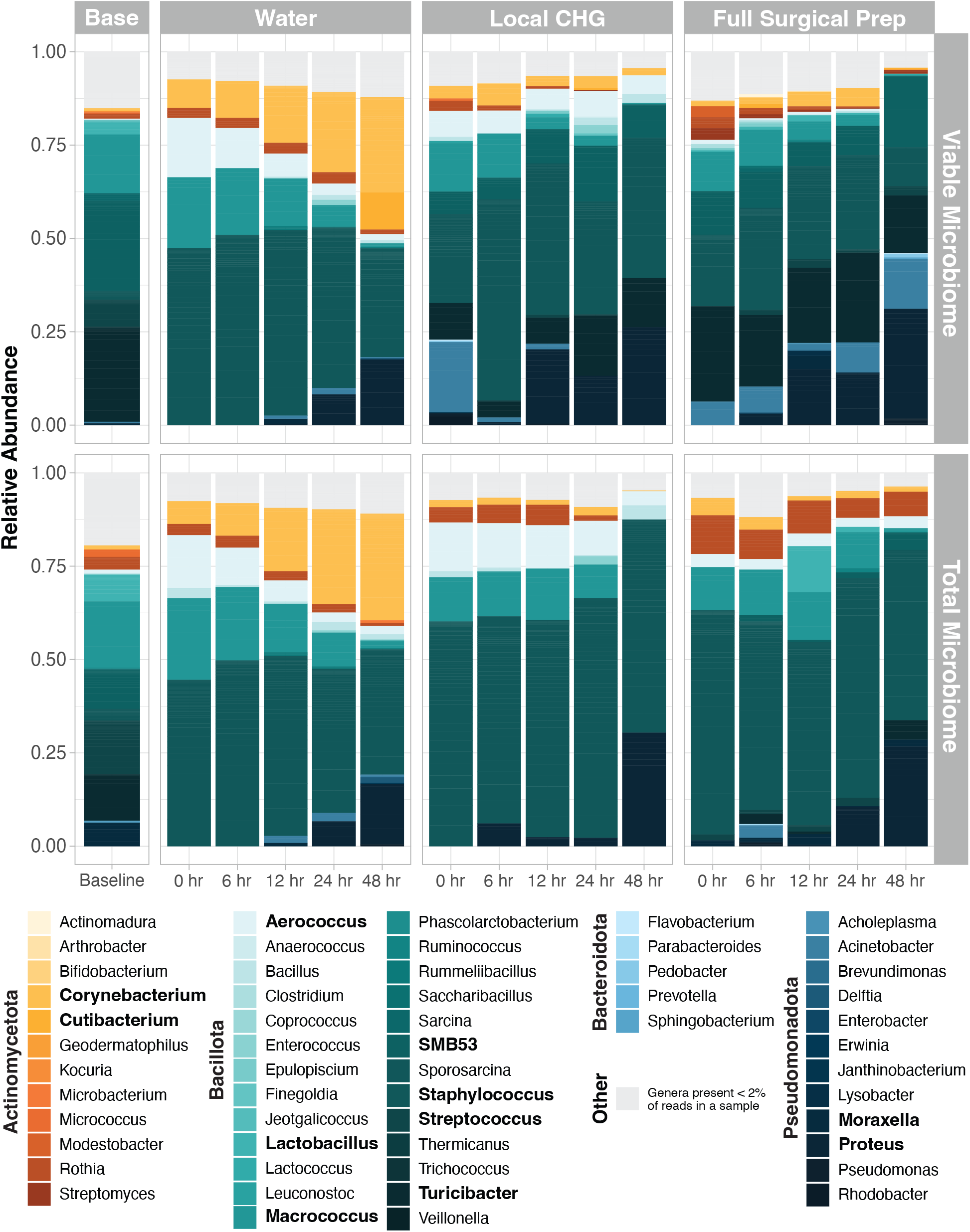
Viable and total *ex vivo* porcine skin microbial communities over time. Plots display the average relative abundance of each genus in the water (control), local CHG application, and full surgical preparation experimental groups. Average genera relative abundance within the viable and total (live + dead) microbial communities are shown in the top and bottom row, respectively. Bolded genera are those comprising at least 30% of the microbial community in at least one sample. Taxa present < 2% in a sample were grouped into the “Other” category.

No significant differences were observed in viable or total microbial community structure in samples collected at baseline or from water treated skin (Multivariate PERMANOVA p-values > 0.5, **Fig. S3A-B, Table S5**). This confirms that in normal homeostatic conditions our viability assay effectively and selectively evaluates DNA from live microorganisms without over representing taxa from a particular phylum or genus. Viable microbiomes are, however, significantly different than the total microbiome following CHG application in both experimental groups (all p-values < 0.001; **Fig. S3C-D, Table S5**), with total microbial communities displaying an over representation of DNA from *Staphylococcus, Macrococcus, Aerococcus*, and *Rothia* (**Fig. S3E-K**). This supports that following antiseptic application free DNA from newly killed bacteria, particularly from highly abundant skin taxa, can persist on the skin and confound sequencing based assessments if viability assays are not appropriately considered.

### Impact of the *Ex vivo* environment on skin microbial communities

Changes in taxonomic proportions over time suggest that the *ex vivo* laboratory environment promotes shifts in the skin microbiome (**Fig. 2, Fig S4**). Compared to baseline, water treated skin undergoes a reduction in the relative abundance of several genera and a corresponding rise in *Staphylococcus* in the first post-treatment timepoint (all FDR q-values < 0.05 **Fig. S4A-H)**. The two CHG treated groups similarly displayed an increase in *Staphylococcus* relative abundance immediately post intervention compared to baseline (all FDR q-values < 0.1; **Fig. S4E and S5A-B**). Over the course of the experiment, all groups experienced significant declines in *Macrococcus* and an increase in *Proteus* (FDR q-value < 0.05; **Fig. S4I-L and S5C-F**). Collectively, these findings highlight that some of viable microbial community changes in the CHG exposed groups are partially secondary to the *ex vivo* laboratory environment.

### Chlorhexidine Gluconate shifts viable skin microbiome structure

The longitudinal experimental design served to characterize the immediate and short-term effects of CHG antiseptic on *ex vivo* skin viable microbial communities. After accounting for skin donor, the viable microbiome after local CHG application, full surgical preparation, and water control groups were significantly different from one another at all timepoints (all p-values < 0.01, multivariate PERMANOVA; **Table S6, Fig 3A**). Samples from both CHG treated groups also displayed significantly higher variance in microbial composition over the course of the experiment, as indicated by larger average distances from their groups’ centroid compared to samples from the water control group (**Fig. 3A, Fig. S6A, Table S7)**. Post-intervention, both CHG treated groups were enriched for *SMB53, Turicibacter, Pseudomonas* and *Proteus* compared to viable microbial communities from water treated skin (all FDR q-value < 0.05; **Fig. 3C-F and Fig. S6B-I, Table S8**). Skin receiving the full pre-surgical preparation had the largest reduction in *Staphylococcus*, the most dominant commensal genera on skin (all FDR q-value <0.0001; **Fig. 3D and Fig. S6I, Table S8**).

**Figure 3:**
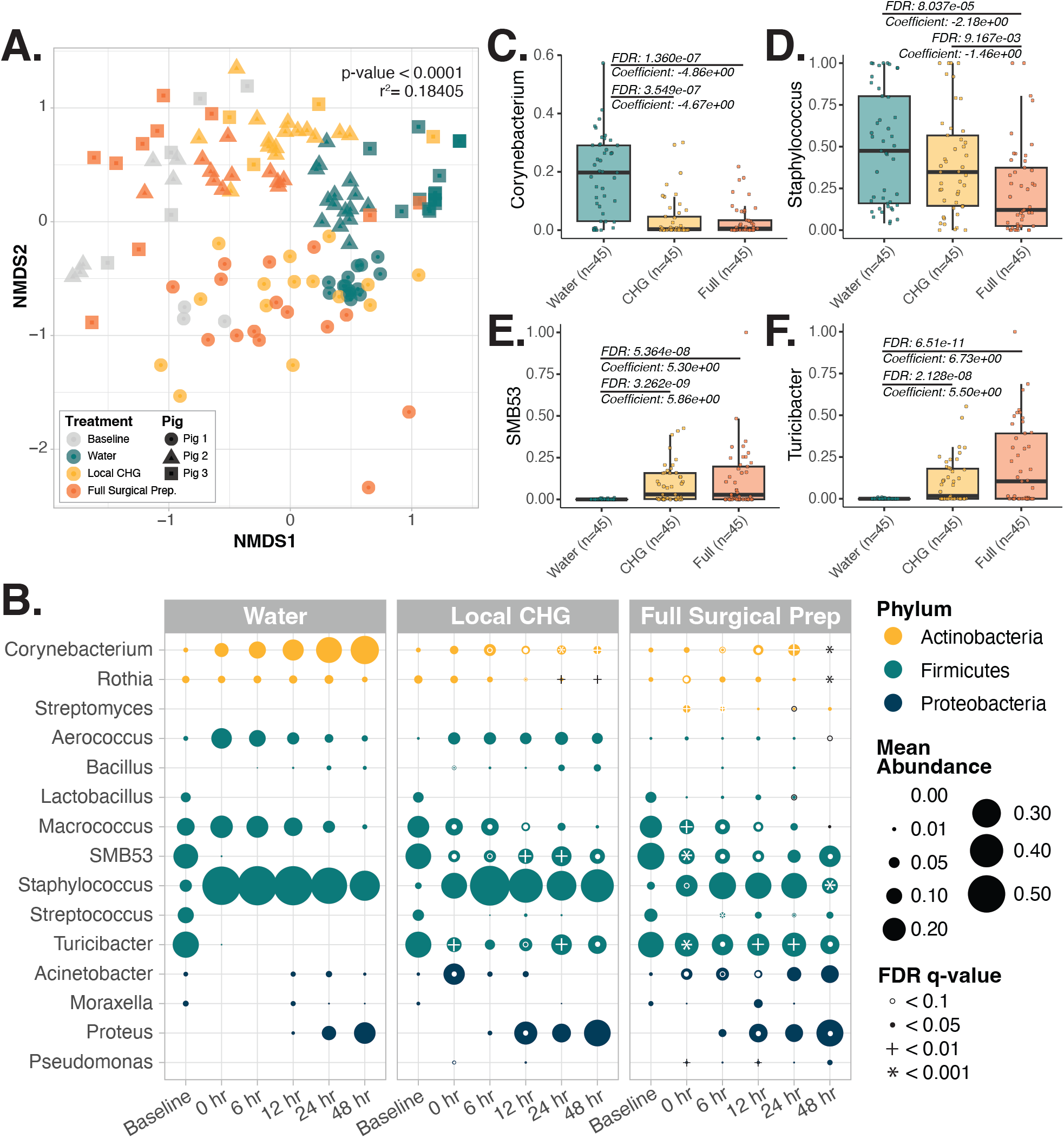
Application of CHG is associated with altered and more variable viable microbial community composition on *ex vivo* skin. **A**. Bray-Curtis beta diversity non-metric multidimensional scaling (NMDS) ordination displaying the variability in sample viable microbial community composition in samples collected at baseline or from skin treated with water or CHG at all post intervention timepoints (0 – 48 hours). Difference in microbial community composition between these groups was evaluated via multivariate PERMANOVA (**Table S6**). **B**. Plot displaying the change in the average relative abundance of key taxa in the viable microbiome on *ex vivo* porcine skin treated with water, a single local CHG application, or the full surgical CHG antiseptic preparation over the course of the experiment. Average taxa abundance across samples is indicated by the size of the point. Differential relative abundance of taxa in the CHG treatment groups to the water treatment group at each respective timepoint was evaluated via MAASLIN2 accounting for the porcine donor as a random effect. White, (or in a few cases black) circles, filled-in dots, plus sign, and asterisk indicate the degree of significance. **C-F**. Plots illustrating significant differences in the relative abundance of key taxa in either of the CHG treatment groups compared to the water control group. MAASLIN2 was used to determine differences in the relative abundance of individual taxa between CHG treated skin compared to water treated skin. These assessments incorporate samples from all post intervention timepoints (0-48 hours). All MAASLIN2 analysis incorporate porcine donor as a random effect and FDR q-values calculated with the Benjamini-Hochberg correction are displayed. Additional plots are in **Supplemental Figure 6**.

Both CHG treatment groups displayed significantly reduced abundance of commensal *Kocuria* (both FDR q-value < 0.05 at 0hr vs. baseline) and increased *Micrococcus* at the first timepoint (both FDR q-value < 0.1 at 0hr vs. baseline; **Fig. 3B and S7-8**). Over the course 48 hours skin microbiomes from both CHG treatment groups saw significant decline in *Rothia (*FDR q-value < 0.01; **Fig. 3B and S7-8**). The proportion of *Proteus* rose in all groups, however this rise began up to 18 hours earlier on CHG treated skin (between 6-12 hours vs. 24-48 hours in the control group), suggesting that the loss of antiseptic induced loss of commensal taxa may allow opportunistic pathogens to dominate sooner (**Fig. 3B and S5E-F**).

### *Ex vivo* skin undergoes lipid remodeling over time

Epidermal and sebaceous lipids serve as key nutrients to skin resident microbial taxa.^42–44^ To explore the influence of the *ex vivo* environment and CHG exposure on epidermal and sebaceous lipids, epidermal punch biopsies were collected at baseline as well as 0- and 48-hours following intervention (**Fig. 1A**). Mass spectrometry based lipidomic assessment identified 1519 lipid species across all samples (**Table S9**). Similar to previous lipidomic characterization of porcine and human skin,^13,45^ ceramides (Cer) and triacylglicerides (TG) comprised the largest proportion of lipids on *ex vivo* porcine skin in terms of abundance (54 +/-13% and 38 +/-13%, respectively, for all samples; **Fig. 4A, Table S9)**. All other groups of lipids, including fatty acids, glycerophospholipids (e.g. phosphatidylcholine), and sterol lipids, collectively comprised less than 10% of the relative abundance (**Fig. 4A, Table S9**). Lipid composition was not associated with either porcine subject or the treatment group, even after accounting for the timepoint of sample collections, suggesting that application of CHG has minimal to no impact on skin lipids (**Fig. S9A-B, Table S10**). Instead, overall lipid composition was significantly associated with the timepoint of sample collection (univariate and multivariate PERMANOVAs < 0.0001, **Fig. 4B, Table S10**). Over time the proportion of ceramides increased from 47 ±8% to 63 ±14% (**Fig. 4A, Table S9**) while triacylglycerides decrease from 44 ± 8 to 31 ± 14 % at baseline and 48 hr time points respectively. The proportion of lipids from other classes decline from 9.4 to 6.3%. Baseline samples displayed significantly greater proportions of several glycerolipids and glycerophospholipids compared to both 0 and 48 hours post intervention (all adjusted Limma p-values < 0.01, **Fig. 4C-F, Table S11**). Meanwhile, samples collected at 48 hours post-intervention had significantly higher proportions of several fatty acids (e.g. fatty acid [FA] 19:0 and FA 20:3) and ceramides (e.g. non-hydroxy-fatty acid sphingosine ceramide [Cer NS] d21:1 30:0 and omega-hydroxy-fatty acid sphingosine ceramide [Cer EOS] d18:1 30:0 18:2) compared to baseline (all adjusted Limma p-values < 0.01, **Fig. 4C-F, Table S11**). Overall, these findings support that shifts in the epidermal and sebaceous lipid composition likely occur secondary to the removal of the skin from the animal and/or the laboratory environment.

**Figure 4:**
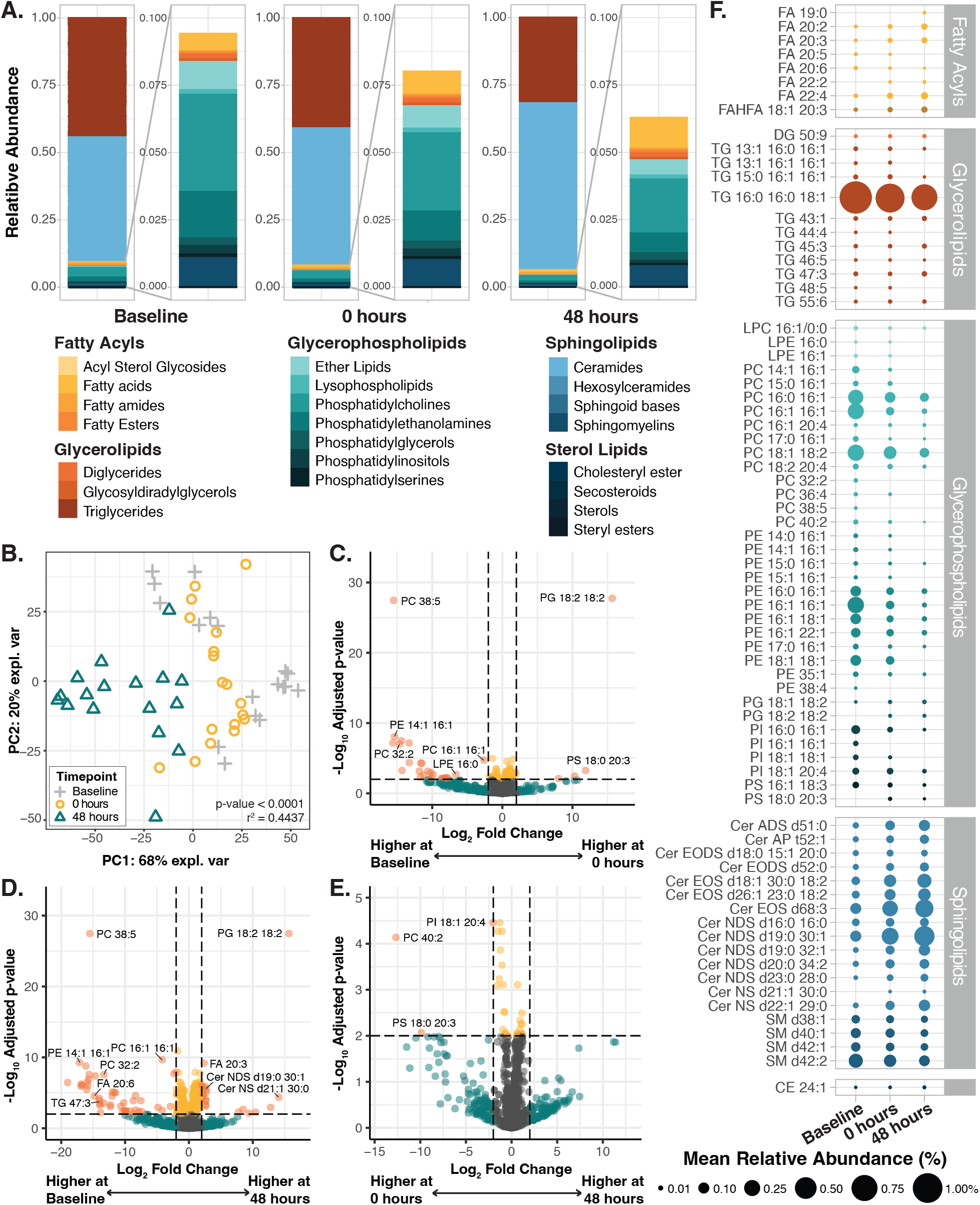
Epidermal and sebaceous lipid composition shifts over the duration of the experiment. Epidermal punch biopsies were collected at baseline as well as 0-and 48-hours following intervention for evaluation of their epidermal and sebaceous lipids **A**. Relative abundance of lipid classes at each timepoint **B**. Principle Coordinate Analysis (PCA) was performed to explore the variability in sample lipid composition. Plot displays the PCA ordination of samples to highlight the association between the timepoint of sample collection and lipid composition. Difference in lipid composition between samples collected at each timepoint was evaluated via univariate PERMANOVA (**Table S10**). Corresponding PCA plots highlighting the associations of lipid composition with porcine subject or experimental group are in **Supplemental Figure 9. C-E**. Volcano plots displaying lipids in significantly higher or lower abundance in samples collected immediately after intervention (0 hr) vs. baseline (**C**), 48 hours after intervention vs. baseline (**D**), or 0 hours vs. 48 hours post intervention (**E**). differential abundance of lipids in samples from each timepoint were evaluated via DEqMS. Lipids were considered differentially abundant in a given group if they displayed log_2_(fold change) > 2 and adjusted Limma p-value < 0.01. **F**. Plot displaying the change in the average relative abundance of key lipid species on *ex vivo* porcine skin at each timepoint. Only lipids identified via DEqMS to be significantly more or less abundant in samples from one of the timepoints are included. Size of the dot indicates the average percent relative abundance. Full results are in **Supplemental Table 11**.

## Discussion

Incomplete antiseptic efficacy against potentially pathogenic Gram-negative and biofilm forming taxa places some patients at disproportionate risk for developing a surgical site infection.^5^ Laboratory models capable of interrogating the effects of antiseptics on the skin and its complex microbial communities are desperately needed to improve and better tailor antiseptic formulations. Within this work we establish an *ex vivo* porcine skin model to study the effects of topical agents on the skin microbiome. We demonstrate that application of the antiseptic CHG results in a temporary reduction of viable microbial bioburden while promoting enrichment of several potentially pathogenic, Gram-negative taxa. Collectively, this work underscores the utility of the *ex vivo* porcine skin system for interrogating chemical disruptions and subsequent recovery of the skin microbiome.

Longitudinal microbial and lipidomic assessments served to evaluate the similarities of the *ex vivo* porcine skin to that of live humans and pigs as well as the impacts of environment on the *ex vivo* tissue and microbial communities. Prior to application of the antiseptic CHG, viable microbial communities on *ex vivo* porcine skin were dominated by taxa within the *Macrococcus, SMB53, Staphylococcus, Streptococcus*, and *Turicibacter* genera, consistent with skin microbiome reports from live pigs.^17,18,46–48^ Similar to prior reports, 85% of the taxa are also observed on human skin.^17^ In the ex vivo system, porcine skin treated with sterile water underwent microbial community restructuring, gradually becoming dominated by *Corynebacterium* and *Staphylococcus*. This composition mirrors human skin microbiomes observed at moist skin sites (e.g. armpit and groin).^42,43^ Moist human skin sites are characterized by high sweat glad density and tend to have warmer skin surface temperatures, around 36.5 - 37.5°C.^49–51^ Its likely that the community shifts observed over the course of the experiment were largely secondary to the relatively warm (37°C) and moist (90-95% humidity) conditions of the experimental conditions promoting the growth of the skin taxa adapted to moist environments. This underscores the potential adaptability of this model to more accurately mimic specific skin microenvironments through modifying the incubator environment parameters.

Lipidomics showed *ex vivo* porcine skin contains triglycerides, diglycerides, free fatty acids, cholesterol esters, and sterol esters likely derived from sebum,^52^ as well as epidermally derived ceramides, phospholipids, and fatty acids.^45,53^ These lipid profiles reinforce the strong similarity of porcine and human skin lipid compositions.^13,14,45,53,54^ Increased relative abundance of ceramides and reduced phospholipids likely represents continued cornification and desquamation without the normal sheading of “dead skin” over the course of the experiment.^55–57^ Consistent abundance of diacylglycerides, cholesterol esters and sterol esters may suggest continued sebaceous gland sebum production.^52^ Meanwhile, the gradual decline in Triglycerides may reflect increased host and microbial breakdown of Triglycerides into free fatty acids over the experiment.^58–60^ Together these findings reinforce the underlying strengths of an *ex vivo* porcine model for the study and inference of human skin function and the potential effects of topical treatments on the skin.

Through the use of a viability assay to more accurately measure the effects of chlorhexidine gluconate on the skin microbiome^5^ the value and unbioased nature of this protocol is demonstrated through the overlap of viable and total microbiomes at baseline and on water treated skin overtime. Notably, the lack of a difference between the water control and the total microbiome of the CHG treated groups immediately after antiseptic application underscores the importance of using a viability assay. For without the viability assay we would not have been able to accurately characterize the impacts of an antiseptic, antibiotic, or other potentially microbially toxic exposure.

By assessing only viable microbes, we demonstrate that CHG induces an immediate reduction in viable microbial bioburden. However, as observed in surgical patients,^5^ CHG does not completely sterilize the skin microbiome. CHG effectively targeted skin commensals (e.g. *Corynebacterium, Macrococcus*, and *Staphylococcus*) while several gut associated and potentially pathogenic taxa persisted on the skin (e.g. *SMB53, Turicibacter, Proteus* and *Pseudomonas)*.^18,61–64^ *Proteus* and *Pseudomonas* are notable biofilm forming Gram-negative pathogens associated with SSI, with *Pseudomonas aeruginosa* being among the most common causes of surgical site infections.^1,65–67^ These findings are consistent with multiple reports that CHG is less effective against Gram-negative bacteria and microbial biofilm.^68–72^ It’s also possible that CHG does not fully penetrate deeper skin layers and hair follicles, where many skin-associated microbes reside.^68–70,73,74^ The very low proportion of *Proteus* and *Pseudomonas* (<1%) in baseline communities also emphasizes how tolerance and resistance to antiseptics can allow rapid expansion following depletion of the community. Collectively, these findings underscore the importance of the skin microbiome in colonization resistance. The loss of commensal taxa following antiseptic exposure may depletes critical protective function leading to a skin microbiome unable to defend against opportunistic pathogens.^42,75– 79^

We also show that reservoirs of viable microbes do persist following antiseptic exposure and are sufficient for microbial bioburden to slowly return after antiseptic exposure. Contrary to our investigations in human subjects,^5^ we did not observe a return of the ex vivo skin microbiome back baseline microbiome compositions. This is likely because unlike in human surgical patients, the only source of microbial repopulation in the *ex vivo* environment are the microbes that survived the antiseptic.

In summary, *ex vivo* porcine skin tissue is a robust model for studying human skin function and microbial community dynamics.^11,12,14^ Our findings reinforce the similarities of human and porcine skin and their microbiomes and highlight the models potential flexibility to mimic specific skin microenvironments. With this model we demonstrate that application of CHG antiseptic temporarily reduces viable microbial bioburden, yet selects for Gram-negative and biofilm forming taxa, which can lead to communities dominated by potential pathogens. Collectively these findings highlight the utility of the *ex vivo* porcine skin system for pre-clinical assessment of topical agents on the skin microbiome and lipid composition and ultimately for the testing and development of improved antiseptic formulations.

## Acknowledgements

We extend considerable thanks to the University of Wisconsin Biotechnology Center for their expert guidance and assistance with lipidomic assessments and microbial sequencing.

We also greatly thank the University of Wisconsin Swine Research Center for their care and dedication to animal subjects and assistance with tissue acquisition.

## Author Contributions

**ECT:** conceptualization, methodology, formal analysis, investigation, data curation, visualization, writing – original draft, writing-review & editing. **KX:** methodology, validation, formal analysis, investigation, data curation, visualization, writing-review & editing. **KDLC:** methodology, investigation, data curation, writing-review & editing. **LH:** methodology, investigation, data curation, writing-review & editing. **SS:** methodology, investigation, data curation, writing-review & editing. **JJM:** resources, writing-review & editing. **DS:** resources, writing-review & editing. **AH:** supervision, resources, writing-review & editing. **AG:** conceptualization, supervision, resources, funding acquisition, writing – original draft, writing-review & editing. **LRK:** conceptualization, supervision, resources, funding acquisition, writing – original draft, writing-review & editing.

## Supplemental Figure Legends

**Supplemental Figure 1: The *ex vivo* porcine skin model. A**. Photo depicting an example of the *ex vivo* porcine skin model. Sections of *ex vivo* skin were cleaned and shaved and cut into roughly 4×7 inch sections for each experimental treatment. Tissue to be used in experiments lasting more than a few hours were placed onto 9×9 inch plates of DMEM agar gel as shown. Each skin section (experimental treatment) was placed on its own separate plate. **B**. Side view of the tissue shown in panel A to highlight the epidermal, dermal, and subcutis skin layers. Note, this particular section of skin is on the thicker side of what is typically used in these experiments (about 1 cm). **C**. Photo illustrating dissection of the epidermis from 12mm punch biopsies for lipidomic assessment.

**Supplemental Figure 2: Determining an optimal PMAxx concentration for selective evaluation of live bacteria within *ex vivo* porcine skin microbial communities**. Four different concentrations of PMAxx were evaluated for accurate quantification of viable and total bacterial bioburden on *ex vivo* porcine skin. Swabs of the skin microbiome were collected; one sample served as a control to determine the anticipated amount of viable bacteria on the skin under normal circumstances, one microbiome sample was intentionally heat killed at 95C for 10 minutes, and one sample was taken from skin following application of CHG antiseptic (n=3 for each group). Bioburden was quantified via viability-qPCR. Percentage of live bacteria was calculated by dividing the number of viable bacteria in the sample by the total number of bacteria. The percent live bacteria in heat killed samples and samples from CHG treated skin were compared to the control group via t-tests with Welch’s correction. * indicates p-value < 0.05

**Supplemental Figure 3: Viable and total microbial community compositions differ only in the CHG treated groups after antiseptic application. A-D**. Non-metric Multidimensional Scaling (NMDS) ordination of the Bray-Curtis dissimilarity matrix for samples collected at baseline (**A**), as well as in the water treated (**B**), local CHG treated (**C**), and full-surgical CHG antiseptic preparation (**D**) groups at all post treatment timepoints (0 through 48 hours post intervention). For each panel multi-variate PERMANOVAs accounting for porcine subject with 9999 permutations were utilized to evaluate the differences between the viable (PMAxx treated; blue) and total (not treated; yellow) sample community compositions. Details can be found in **supplemental table 5. E-K**. MAASLIN2 was used to determine differences in the relative abundance of individual taxa between viable and total communities on *ex vivo* skin treated with CHG. Microbial communities from the local CHG and full surgical prep treatment groups at all post intervention time points (0 through 48 hours) were evaluated together. For all MAASLIN2 analyses, porcine donor was incorporated into the assessment as a random effect. FDR q-values calculated with the Benjamini-Hochberg correction are displayed.

**Supplemental Figure 4: The laboratory environment promotes shifts in the *ex vivo* porcine skin microbial communities. A-H**. MAASLIN2 was used to determine differences in the relative abundance of individual taxa between the baseline and immediately post intervention (0 hr) timepoint within the viable microbial communities on control ex vivo porcine skin treated with only sterile water. FDR q-values calculated with the Benjamini-Hochberg correction are displayed. **I-L**. MAASLIN2 was also utilized to evaluate the correlations of microbial taxa relative abundance on water treated skin over time. Due to the significant community shifts that occurred between baseline and the post intervention (0hr timepoint; **A-H and Fig 3A-B**), only samples from post intervention timepoints were included in this analysis. For all MAASLIN2 analyses, porcine donor was incorporated into the assessment as a random effect. FDR q-values calculated with the Benjamini-Hochberg correction are displayed.

**Supplemental Figure 5: Some changes seen following CHG treatment may in part be secondary to the laboratory environment**. Panels highlight changes within skin microbiome following local CHG or full surgical preparation with CHG that mirror changes observed in the microbial communities on water treated skin (**Fig. S4)**. Panels on the left hand side are changes seen on skin treated with local CHG and panels on the right hand side are changes seen on skin given the full surgical CHG preparation. **A-B** MAASLIN2 was used to determine differences in the relative abundance of individual taxa between the baseline and immediately post intervention (0 hr) timepoint within the viable microbial communities on ex vivo porcine skin for both of the CHG treatment groups. For all MAASLIN2 analyses, porcine donor was incorporated into the assessment as a random effect. FDR q-values calculated with the Benjamini-Hochberg correction are displayed. **C-F**. MAASLIN2 was also utilized to evaluate the correlations of microbial taxa relative abundance on CHG treated skin over time. Due to the significant community shifts that occurred between baseline and the post intervention (0hr timepoint), only samples from post intervention timepoints were included in this analysis. Additional plots displaying changes specific to either CHG treated group over time are in **Supplemental Figures 7-8**

**Supplemental Figure 6: Application of CHG is associated with altered and more variable viable microbial community composition. A**. Companion to **Figure 3A**. Bray-Curtis dissimilarity metric was used to evaluate the similarity of each sample’s microbial community composition. Distances from each group’s centroid were calculated from this dissimilarity matrix. The water, local CHG, and full surgical prep groups include all samples collected at post intervention timepoints. Tukey multiple comparisons of means was then used to determine if the degree of variability within groups were significantly different. **B-I**. Continuation of **Fig 3C-F**. Relative abundance plots illustrating significant differences in key taxa in either of the CHG treatment groups compared to the water control group. MAASLIN2 was used to determine differences in the relative abundance of individual taxa between CHG treated skin compared to water treated skin. These assessments incorporate samples from all post intervention timepoints (0-48 hours). All MAASLIN2 analysis incorporate porcine donor as a random effect and FDR q-values calculated with the Benjamini-Hochberg correction are displayed. Details on significant differences between the treatment groups at each timepoint individually are displayed in **Fig 3B**.

**Supplemental Figure 7: Changes in microbial taxa relative abundance on skin treated with a single application of CHG. A-F**. MAASLIN2 was used to determine differences in the relative abundance of individual taxa between the baseline and immediately post intervention (0 hr) timepoint within the viable microbial communities on *ex vivo* porcine skin treated with the single local application of CHG. For all MAASLIN2 analyses, porcine donor was incorporated into the assessment as a random effect. FDR q-values calculated with the Benjamini-Hochberg correction are displayed. **G-H**. MAASLIN2 was also utilized to evaluate the correlations of microbial taxa relative abundance on the local CHG treated skin over time. Due to the significant community shifts that occurred between baseline and the post intervention (0hr timepoint; **A-F and Fig 3B**), only samples from post intervention timepoints were included in this analysis. Additional plots displaying changes over time within the CHG treated groups that mirror those seen on water treated *ex vivo* skin are in **Supplemental Figure 5**.

**Supplemental Figure 8: Changes in microbial taxa relative abundance on *ex vivo* skin following the full surgical preparation with CHG. A-F**. MAASLIN2 was used to determine differences in the relative abundance of individual taxa between the baseline and immediately post intervention (0 hr) timepoint within the viable microbial communities on *ex vivo* porcine skin treated the full surgical CHG preparation. For all MAASLIN2 analyses, porcine donor was incorporated into the assessment as a random effect. FDR q-values calculated with the Benjamini-Hochberg correction are displayed. **G-L**. MAASLIN2 was also utilized to evaluate the correlations of microbial taxa relative abundance on the full surgical prep treated skin over time. Due to the significant community shifts that occurred between baseline and the post intervention (0hr timepoint; **A-F and Fig 3B**), only samples from post intervention timepoints were included in this analysis. Additional plots displaying changes over time within the CHG treated groups that mirror those seen in on water treated *ex vivo* skin are in **Supplemental Figure 5**.

**Supplemental Figure 9: Epidermal and sebaceous lipid composition is not associated with porcine subject or experimental group. A-B**. Principle Coordinate Analysis (PCA) was performed to explore the variability in sample lipid composition. These plots are companion plots to **Figure 4B**. All three plots are the same ordination colored differently to highlight the (lack of) association between porcine subject (A) or experimental treatment group (B) with sample lipid composition. Difference in lipid composition between samples collected at each timepoint was evaluated via univariate PERMANOVA (both p-values > 0.1; **Table S10**).

## Supplemental Table legends

**Supplemental Table 1: Percentage of live bacteria in initial experiments to optimize viability-PCR parameters**. Percent live bacteria represents the calculated viable bacteria divided by the calculated total bacteria. Data represented as mean ± standard deviation (n = the number of samples included from each experiment). N indicates the number or replicates in each group. For each respective PMAxx concentration, heat killed and CHG treated groups were compared to the control group via t-tests with Welch’s correction.

**Supplemental table 2: Viable and total microbial bioburden on *ex vivo* porcine skin treated with water or following CHG antiseptic over time**. Viable and total bacterial bioburden were determined via viability-qPCR. Data displayed as mean ± SD Log10[Bacteria]. Comparisons between bacterial bioburden in the local CHG and full surgical prep groups to the water treated control group at the same timepoint were made via t-tests with Welch’s correction (asterisk). Total and viable bacterial burden within a experimental group at each timepoint were compared via paired t-tests (far right column; open circles and dots). ∘ p-value < 0.1 Total v. Viable at same timepoint (trending toward significance); • p-value < 0.05 Total v. Viable at same timepoint; •• p-value < 0.01 Total v. Viable at same timepoint; * p-value < 0.05 v. water control group at same timepoint; ** p-value < 0.01 v. water control group at same timepoint; *** p-value < 0.001 v. water control group at same timepoint.

**Supplemental Table 3: 85% of bacterial genera identified on *ex vivo* porcine skin have also been found in human skin microbial communities**. References where these bacteria have been noted in human skin microbial communities are noted.

**Supplemental Table 4: Experimental factors associated with microbial community composition**. Bray Curtis beta diversity was utilized to compare sample community compositions across all total and viable microbial community samples. Associations between experimental factors with microbial community compositions were evaluated first with Univariate PERMANOVAS as well as multi-variate PERMANOVAS accounting for the porcine subject. Each PERMANOVA was run considering the marginal effects of terms with 9999 permutations.

**Supplemental Table 5. Viable and total microbial community compositions differ following CHG application**. Univariate PERMANOVAs and multivariate PERMANOVAS with 9999 permutations were utilized to evaluate the differences between the viable (PMAxx treated, containing only DNA from viable bacteria) and total (not treated, containing DNA from both viable and dead bacteria) microbial community compositions on control (water), local CHG treated, or full surgical prep CHG treated skin at each timepoint and all post-intervention timepoints combined. Corresponding NMDS plots are in **supplemental figure 3**.

**Supplemental Table 6. Following CHG application, skin microbiome compositions significantly differ from the water treated control**. Companion to **Figure 3**. Univariate PERMANOVAs and multivariate PERMANOVAS with 9999 permutations were utilized to evaluate the differences between the microbial communities on *ex vivo* skin treated with either a single CHG application, the full surgical preparation, or just sterile water. Groups were compared both at each timepoint separately and all post intervention timepoints collectively.

**Supplemental Table 7: Compared to water treated skin, skin microbial communities display greater variability in composition following CHG antiseptic application**. Companion to **Figure 3A and Figure S6A**. Bray-Curtis dissimilarity metric was used to evaluate the similarity of each sample’s microbial community composition. Distances from each group’s centroid were calculated from this dissimilarity matrix. The water, local CHG, and full surgical prep groups include all samples collected at post intervention timepoints. Tukey multiple comparisons of means was then used to determine if the degree of variability within groups were significantly different.

**Supplemental Table 8: Significant differences in taxa relative abundance on *ex vivo* skin treated with CHG compared to skin treated with sterile water at all post intervention timepoints**. This table is a companion to **Figure 3 and Supplemental Figure 6**. MAASLIN 2 was used to evaluate differential relative abundance of taxa within viable microbial communities on *ex vivo* skin following either a single application of CHG or following the series of CHG applications for full surgical preparation. Analyses included samples from each experimental group at all post intervention timepoints (0-48 hrs). All analyses were conducted incorporating the porcine donor as a random effect.

**Supplemental Table 9: Major classes of lipids in and on *ex vivo* porcine skin epidermis**. Table includes the number of lipid species within each class identified across all epidermal biopsy samples. Subsequent columns display the relative abundance (mean ± standard deviation) of each lipid class in all samples or from samples collected at each timepoint.

**Supplemental Table 10: Timepont of epidermal biopsy collection has the strongest association with sample lipid composition**. Companion to **Figure 4B** and **Supplemental Figure 9**. Univariate PERMANOVAs and multivariate PERMANOVAS with 9999 permutations were utilized to evaluate for potential differences in sample epidermal and sebaceous lipid composition between various experimental factors.

**Supplemental Table 11: The abundance of several lipid species significantly changes over the duration of the experiment**. Companion to **Figure 4C-E**. Differential abundance of lipids in samples from each timepoint were evaluated via DEqMS. Only lipids considered differentially abundant in a given group are included. Lipids were considered significant if they displayed log_2_(fold change) > 2 and adjusted Limma p-value < 0.01.

